# Ultrafast amplitude modulation for molecular and hemodynamic ultrasound imaging

**DOI:** 10.1101/2021.05.18.444561

**Authors:** Claire Rabut, Di Wu, Bill Ling, Zhiyang Jin, Dina Malounda, Mikhail G. Shapiro

## Abstract

Ultrasound is playing an emerging role in molecular and cellular imaging thanks to new micro- and nanoscale contrast agents and reporter genes. Acoustic methods for the selective in vivo detection of these imaging agents are needed to maximize their impact in biology and medicine. Existing ultrasound pulse sequences use the nonlinearity in contrast agents’ response to acoustic pressure to distinguish them from mostly linear tissue scattering. However, such pulse sequences typically scan the sample using focused transmissions, resulting in a limited frame rate and restricted field of view. Meanwhile, existing wide-field scanning techniques based on plane wave transmissions suffer from limited sensitivity or nonlinear artifacts. To overcome these limitations, we introduce an ultrafast nonlinear imaging modality combining amplitude-modulated pulses, multiplane wave transmissions and selective coherent compounding. This technique achieves contrast imaging sensitivity comparable to much slower gold-standard amplitude modulation sequences and enables the acquisition of larger and deeper fields of view, while providing a much faster imaging framerate of 3.2kHz. Additionally, it enables simultaneous nonlinear and linear image formation, and allows concurrent monitoring of phenomena accessible only at ultrafast framerates, such as blood volume variations. We demonstrate the performance of this ultrafast amplitude modulation (uAM) technique by imaging gas vesicles, an emerging class of genetically encodable biomolecular contrast agents, in several in vitro and in vivo contexts. These demonstrations include the rapid discrimination of moving contrast agents and the real-time monitoring of phagolysosomal function in the mouse liver.

## INTRODUCTION

Ultrasound imaging enables the assessment of organ anatomy and function with high spatial and temporal resolution (typically < 500 μm and 10 ms). Recently, ultrasound has gained increasing capabilities for molecular and cellular imaging due to the development of micro- and nanoscale contrast agents and reporter genes capable of targeting specific disease states^1–3^ or visualizing cellular processes such as gene expression and enzyme activity^4–6^. For example, biomolecular contrast agents known as gas vesicles (GVs)^7^ can be used to enhance hemodynamic imaging^8^, visualize lysosomal function^9^, functionalized with binding domains to target specific cells^10^, or expressed heterologously as reporter genes or biosensors in bacteria^4–6^ and mammalian cells^5^. GVs comprise a 2-nm-thick protein shell, enclosing cylindrical compartments of air with a typical diameter of ~85 nm and length of 500 nm^11^. Certain GV types exhibit strongly nonlinear responses to acoustic pressure, manifesting as reversible buckling of their shell^12–14^, which results in nonlinear scattering of ultrasound (**Fig.1 a-b**). Contrast agents such as GVs can be detected with improved sensitivity and specificity using ultrasound imaging paradigms such as amplitude-modulation (AM). Parabolic amplitude modulation (pAM)^13^ and cross-propagating amplitude modulation (xAM)^15^ pulse sequences perform line-by-line scans of the media, transmitting triplets of pulses of relative amplitudes ½, ½ and 1. Despite being well-suited for GV imaging in biological samples, both pAM and xAM are limited by their imaging depth and framerate, preventing the monitoring of fast nonlinear events across the imaging plane. Ideally, nonlinear imaging of contrast agents should cover the entire field of interest, provide deep penetration, produce a fast framerate and allow the simultaneous acquisition of other information such as blood flow or tissue motion.

**Figure 1 :**
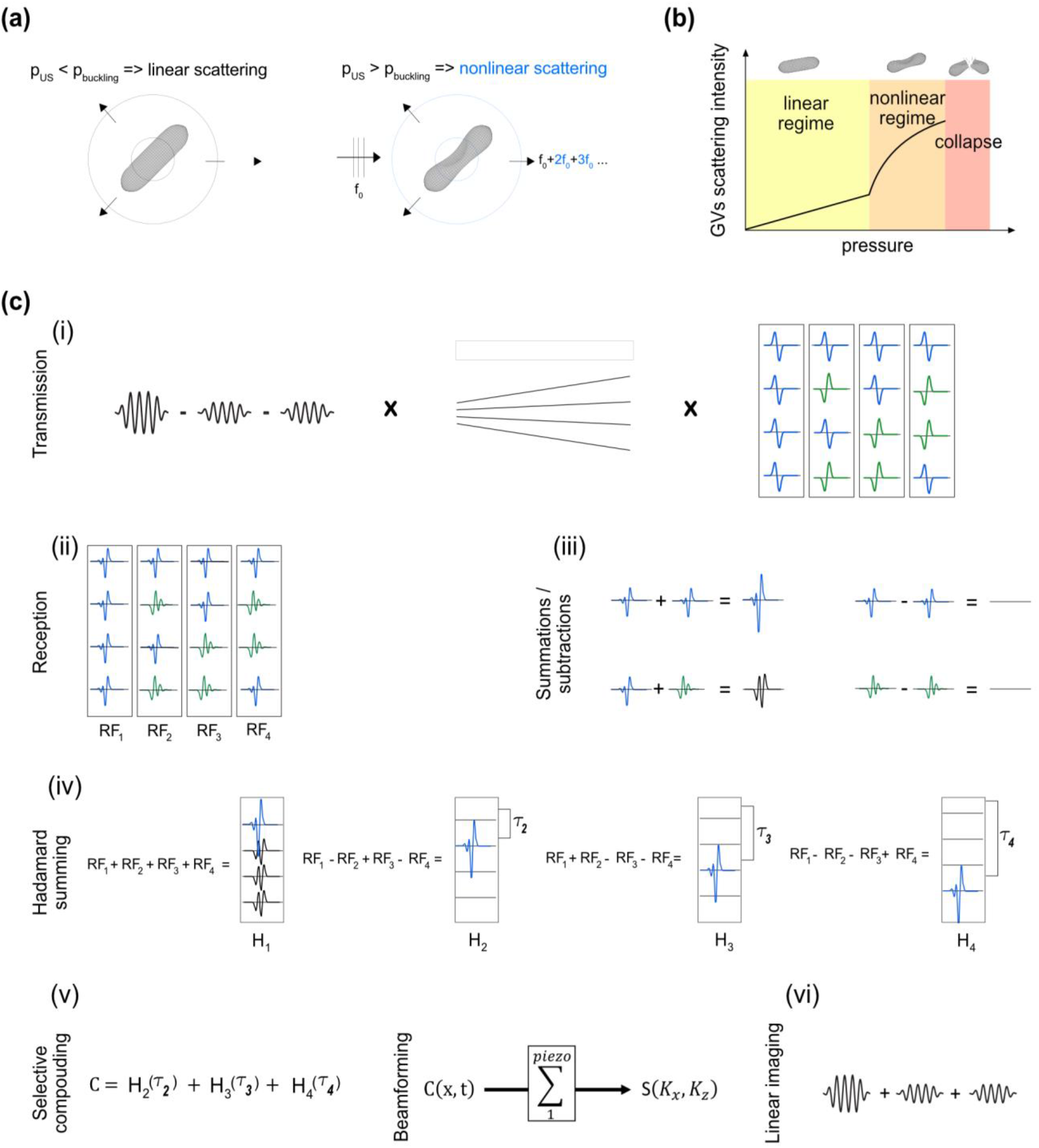
Nonlinear response of GVs and schematic representation of the uAM pulse sequence with N=4 angles. (a) The nonlinear scattering behavior of GVs insonified above their buckling pressure enables their detection with AM (adapted from 15). (b) Schematic scattering intensity curve of GVs as a function of incident pressure. (c) Schematic representation of the uAM pulse sequence for N=4 angles: (i) AM pulses are combined with MPW transmissions^17^ and repeated four times with different polarizations. The polarity combinations are given by the columns of the Hadamard matrix of order 4. (ii) After subtraction of the signals elicited by the two half-amplitude pulses from the full-amplitude one, the nonlinear response appears in the RF data. (iii) The “Hadamard summing” step results in the following combinations: addition of positive nonlinear echoes, subtraction of positive nonlinear echoes, addition of opposite polarity nonlinear echoes and subtraction of negative nonlinear echoes. The results of these combinations are respectively: a positive echo of amplitude N (N=number of summed echoes), zero, a nonlinear signal, zero. (iv) Following the lines of the Hadamard matrix of order 4, “Hadamard RF” data are obtained where each plane wave is retrieved individually with an amplitude 4. However, H1 also exhibits successive nonlinear echoes. (v) To obtain a final image without nonlinear artifacts, coherent compounding is applied with all the Hadamard RF except H1, and a last beamforming step creates a nonlinear image. (vi) The same acquisition can be used for linear monitoring by summing the contributions of half-amplitude pulses instead of subtracting them.

To address these challenges, we introduce ultrafast amplitude modulation (uAM), a nonlinear paradigm inspired by coherent plane wave compounding for very high frame rate ultrasonography^16^. uAM acquires nonlinear images through the coherent summation of ultrasound signals obtained after transmission of successive tilted, amplitude modulated, plane waves. In general, plane wave imaging enables rapid one-shot coverage of the entire field but generates less signal relative to noise per pulse than parabolic or cross-propagating paradigms. The three typical solutions to counterbalance this limitation are increasing the transmit amplitude, increasing the number of compounded angles or averaging multiple sequentially acquired images. However, contrast agents such as GVs require imaging within a specific range of pressures (**Fig.1.b**) – typically between 200 kPa (nonlinear regime) and 600kPa (collapse) for the GV type used in this study^10^, while temporal averaging – by increasing the number of tilted angles or averaging multiple sequential images – leads to a reduced framerate. To overcome these limitations, uAM combines amplitude modulation with multiplane-wave (MPW) transmission and selective coherent compounding. MPW imaging, which comprises the successive transmission of polarized plane waves within a single echo return period, was introduced in 2015 as a means to increase the linear signal-to-noise ratio (SNR) in ultrafast imaging^17^. We endeavored to perform MPW nonlinear imaging by using MPW transmissions with modulated amplitudes. However, because it is based on the recombination of inverted polarity pulses, the Hadamard encoding scheme of MPW transmissions generally requires the assumption of linear scattering, with nonlinear media producing pulse inversion artifacts^18,19^. We hypothesized that we could circumvent this limitation by introducing a new coherent compounding approach that takes selective advantage of those Hadamard sums in which the pulse inversion artifacts are cancelled. Moreover, we hypothesized that the ultrafast frames recorded during the resulting AM-adapted MPW acquisitions could be processed for dynamic linear and nonlinear signals, enabling, for example, simultaneous blood flow measurement^20^. In this Letter, we start by describing the ultrasound transmission sequence and signal processing algorithm underlying uAM. Next, we compare the performance of uAM to pAM and xAM in static nonlinear imaging of GVs. We then test the ability of uAM to visualize the motion of flowing GVs in acquisitions lasting a few milliseconds. Finally, we use uAM to image phagolysosomal function in mice, demonstrating the ability to visualize both biomolecular contrast and blood flow simultaneously in a single pulse sequence.

## RESULTS

Bursts of N successive tilted plane waves are repeated three times with modulated amplitude: two bursts of half amplitude (achieved by silencing respectively the odd and even elements of the transducer) and one burst of full amplitude. After reception, the subtraction of the two half-amplitude bursts from the full amplitude burst allows the elimination of linear signal and the capture of specifically nonlinear responses (**Fig.1.c.ii**). On the contrary, the addition of the modulated pulses results in the capture of the linear response. The trios of modulated MPW bursts are then repeated N times, and for each repetition the polarities of the successive plane waves are given by the column of the Hadamard matrix of order N (**Fig.1.c.i**, **Appendix**).

In conventional MPW imaging, the contribution of all the Hadamard RF data is summed using appropriate offsets of *τ_i_* to produce a coherent recombination of compounded plane wave signals. We however found that the first Hadamard summation systematically leads to a pulse-inversion like summation of polarized pulses^21^ and therefore carries inappropriately time-delayed echoes in the recombined signals. This problem is created by nonlinear media such as contrast agent inclusions and has been described in ^18,19,22^. In addition, we found that all subsequent Hadamard summations do not result in pulse inversion artifacts (**Fig.1.c.ii-iv**, **Appendix**). To therefore benefit from the strength of MPW imaging without creating undesirable nonlinear artifacts due to recombined polarized signals, we introduce selective coherent compounding of all Hadamard RF data except the first one (H_1_), obtaining a compounded data set C (**Fig.1.c.v**):

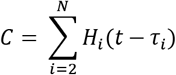

We then can beamform C by applying a delay-and-sum algorithm and obtain a final nonlinear image S, where harmonic residues at inappropriate time delays resulting from pulse inversion nonlinearity have been eliminated. The resulting image is expected to contain (N-1) times the signal obtained with a single plane wave. Linear reconstruction similarly benefits from selective compounding with a reduction of nonlinear residues (**Fig.1.c.vi**).

To validate the uAM pulse sequence, we first imaged a 200 μm-diameter inclusion of GVs (1 nM) in a 1% agarose phantom (**Fig.2.a**) using 4 angles (uAM-4) (**Fig.2.b**). Subtraction of the signals elicited by the two half-amplitude bursts from those received after full-amplitude transmission resulted in backscattered AM echoes along the vertical line crossing the inclusion after each MPW burst (**Fig.2.c**). After standard Hadamard summation, the sums H_2_, H_3_ and H_4_ each exhibit a single high amplitude echo with an appropriate time delay, but H_1_ carries four echoes: the first resulting from the summation of the four in-phase backscattered echoes, and three others resulting from the summing of nonlinear echoes of opposite polarities (**Fig.2.d**). As a result of this nonlinear artifact, AM images generated by conventional compounding of the Hadamard-RF data show three successive nonlinear residues (**Fig.2.e**). On the other hand, selective compounding of all the Hadamard sums other than H_1_ results in an image showing only the correct GV inclusion (**Fig.2.f**). These results confirm that the selective compounding approach enables uAM to take advantage of the MPW paradigm without nonlinear Hadamard summation artifacts.

**Figure 2:**
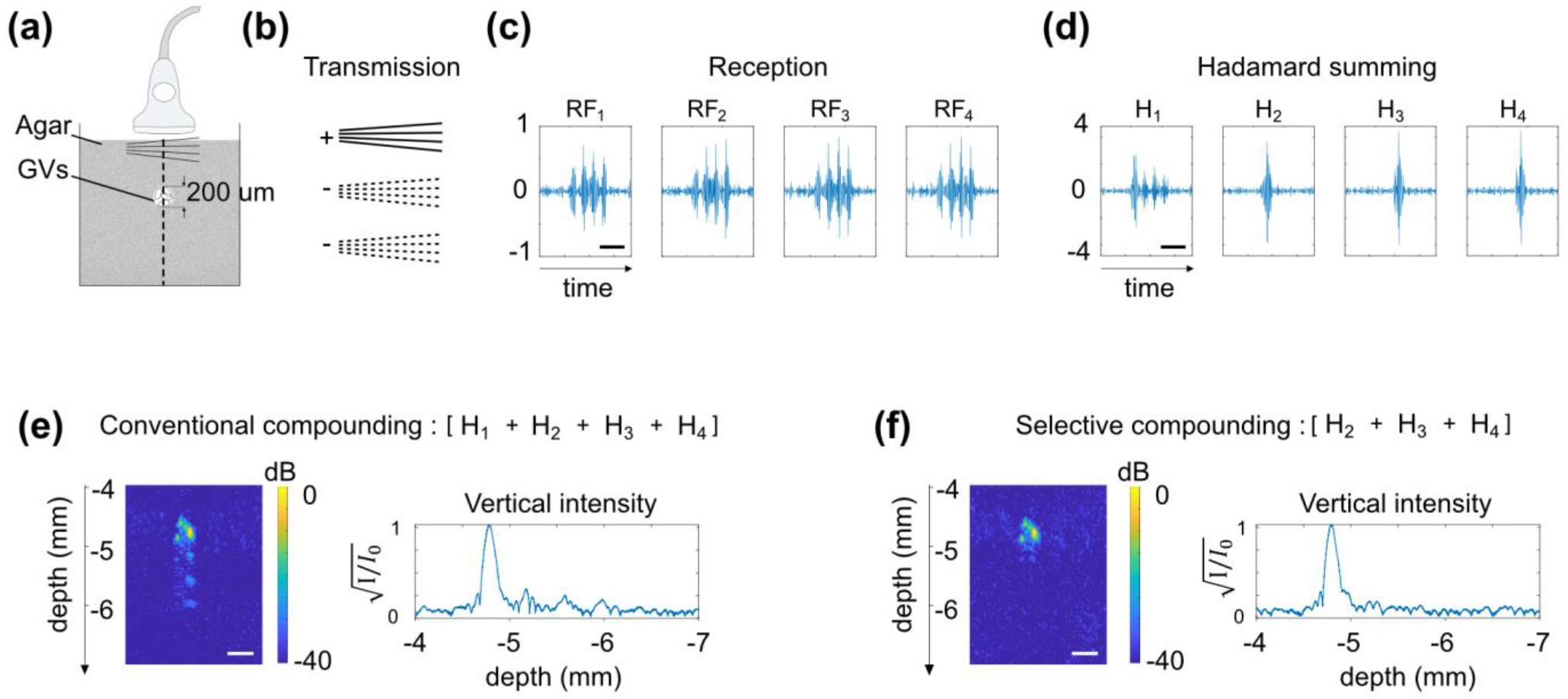
Selective compounding eliminates the nonlinear artifacts of Hadamard summation. (a) Schematic of an agarose phantom in which a 200 μm diameter inclusion of GVs (1000 pM) is embedded. (b) Transmission sequence and signs of the summation at reception of the AM bursts (N=4 angles). (c) RF data plotted along the vertical line crossing the inclusion. (d) After Hadamard summing, individual plane waves are retrieved with amplitude N, except for H1, which contains additional nonlinear echoes. (e) Conventional coherent compounding integrates the artifactual nonlinear residues into the final image. (f) Selective compounding omitting H1 enables the reconstruction of a nonlinear image sans artifact. Time scale bars (c.d.): 1us. Lateral dimension scale bars (e.f.): 100um. Depths calculated from the surface of the probe.

After establishing its basic functionality, we evaluated the performance of uAM with varying angle number (**Fig.3.a**) in comparison to pAM^13^ and xAM^15^ (**Fig.3.b,c**) in a tissue-mimicking phantom containing two rows of 2mm diameter wells filled with GVs. We quantified the contrast-to-tissue ratio for both the upper and deeper inclusions 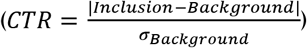, as well as the contrast-to-artifact ratio 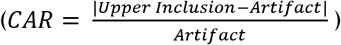.

**Figure 3:**
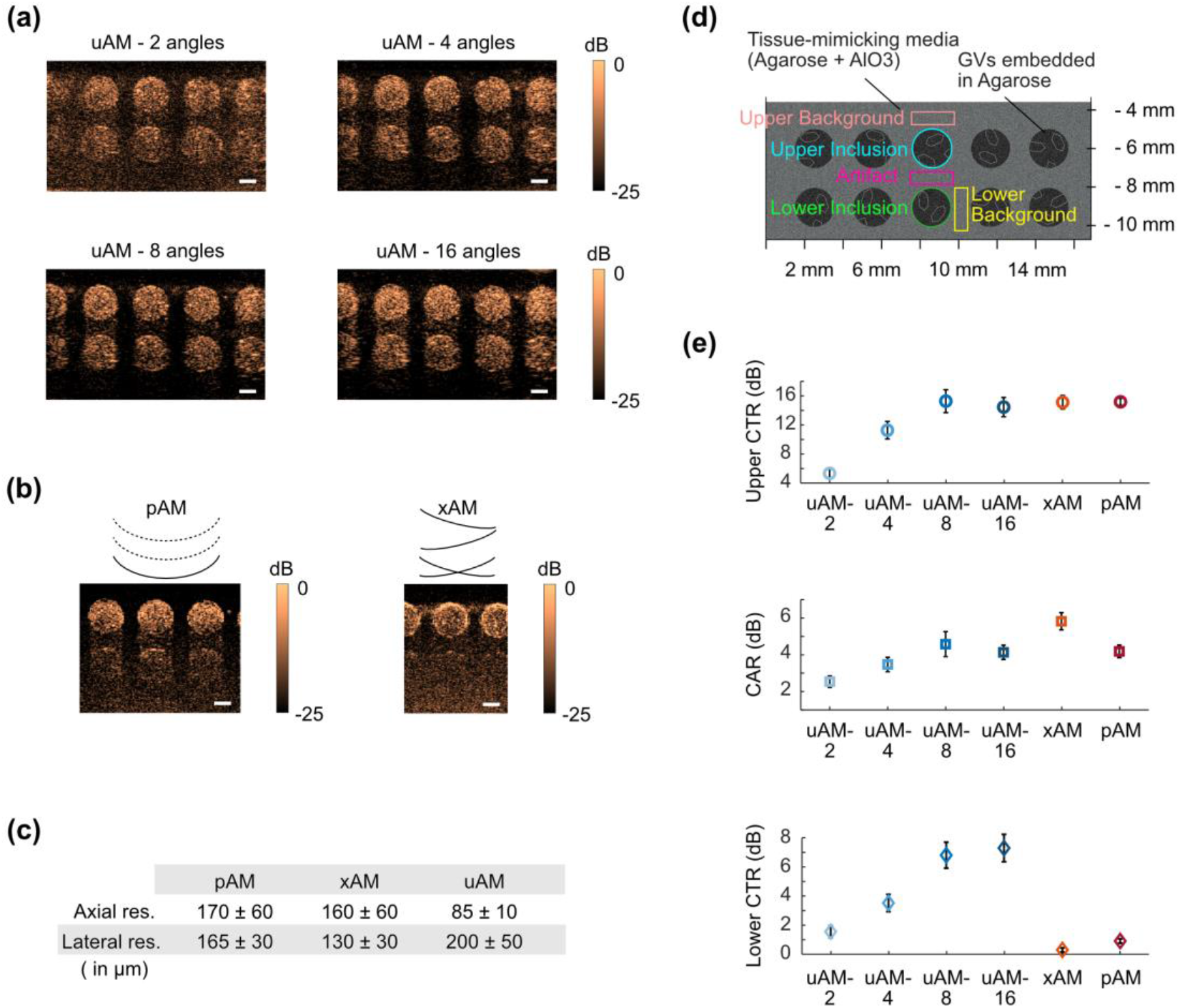
In vitro evaluation of uAM performance in comparison with pAM and xAM. (a) uAM images of GVs embedded in a tissue-mimicking phantom as a function of the number of tilted angles. (b) Same imaging plane as in (a) acquired with pAM and xAM. (c) Spatial resolution of pAM, xAM and uAM calculated with an agar phantom containing sub-wavelength lines of GVs. ± represents STD (d) Schematic of the tissue-mimicking phantom. Quantification was performed on the 3 middle inclusions of each image over 3 different sample replicates. (e). Upper CTR, CAR and Lower CTR for each pulse sequence. The Upper CTR is evaluated between the upper inclusions and the upper background. The CAR is evaluated between the upper inclusions and the artifact region underneath. The Lower CTR is calculated between the lower inclusion and the lower background. Error bars represent ± SEM for N=3 sample replicates with 3 wells per replicate. Scale bars: 1 mm.

We immediately observe that uAM offers the widest and deepest field of view compared to pAM and xAM. uAM scans the entire imaging plane below the transducer array with each transmission. In contrast, the lateral and depth coverage of pAM are limited by transmission aperture and axial focusing, while xAM is limited by the aperture requirements of cross-propagating waves and the maximal wave intersection depth^15^. In terms of image quality (**Fig.3.e**), the highest CTR values for the upper sample were obtained with xAM, pAM and uAM-8 and −16 (all around CNR = 15), decreasing substantially for uAM with smaller numbers of angles. The CAR, a measure of resilience to nonlinear propagation artifacts, was best with xAM, which was specifically developed to eliminate such artifacts through cross-propagation^15^. uAM-8 and −16 angles also provided improved CAR compared to pAM. Finally, the most striking differences in CTR concerned the lower inclusions, which are barely detectable in pAM and xAM, but are robustly seen with uAM, albeit with a lower CTR than the upper row due to attenuation. These results allow us to conclude that uAM with N_angles≥8 offers wider and deeper fields of view than pAM and xAM imaging, comparable near-field CNR, superior CTR at depth, and CAR performance between those of pAM and xAM. 8-angle uAM was selected as the pulse sequence for the remainder of our study.

To compare the lateral and axial resolution of the three AM sequences, we imaged an agar phantom containing sub-wavelength lines of GVs (**Fig.S3**). This experiment revealed that the lateral resolution in uAM is slightly diminished compared to the classic methods (17% larger than in pAM), while axial resolution is improved (by 50% compared to pAM) (**Fig.3.c**).

Remarkably, the performance improvement described above was obtained while accelerating the imaging frame rate by more than one order of magnitude. For an equivalent 10-mm-deep field, acquiring a pAM or xAM image requires approximately 4 ms, while a uAM-8 image is obtained in just 0.31 ms (Appendix). To demonstrate the utility of this acceleration, we evaluated the capacity of uAM to visualize flowing GVs in vitro. Our experimental set-up (**Fig.4.a**) comprised a phantom made of stationary GVs (114 pM) embedded in agarose gel, in which two horizontal tunnels of diameter 1mm allowed the flow of chosen solutions. A flow of 1 mL/min was created by pump-driven syringes. Comparing xAM (**Fig.4.b**) and uAM (**Fig.4.c**), we first imaged the phantom in which only PBS was flowing through the tunnels. As expected, the tunnels within the effective field of view were devoid of contrast. In xAM the lower tunnel was obscured by the depth limitation of the sequence, and a strip of higher signal was also evident. We interpret the latter effect as an artifact of the beamforming algorithm, which assumes a uniform supersonic velocity for the plane wave intersection, while boundary effects in the transmitted waves cause imperfect planarity at their ends.

**Figure 4:**
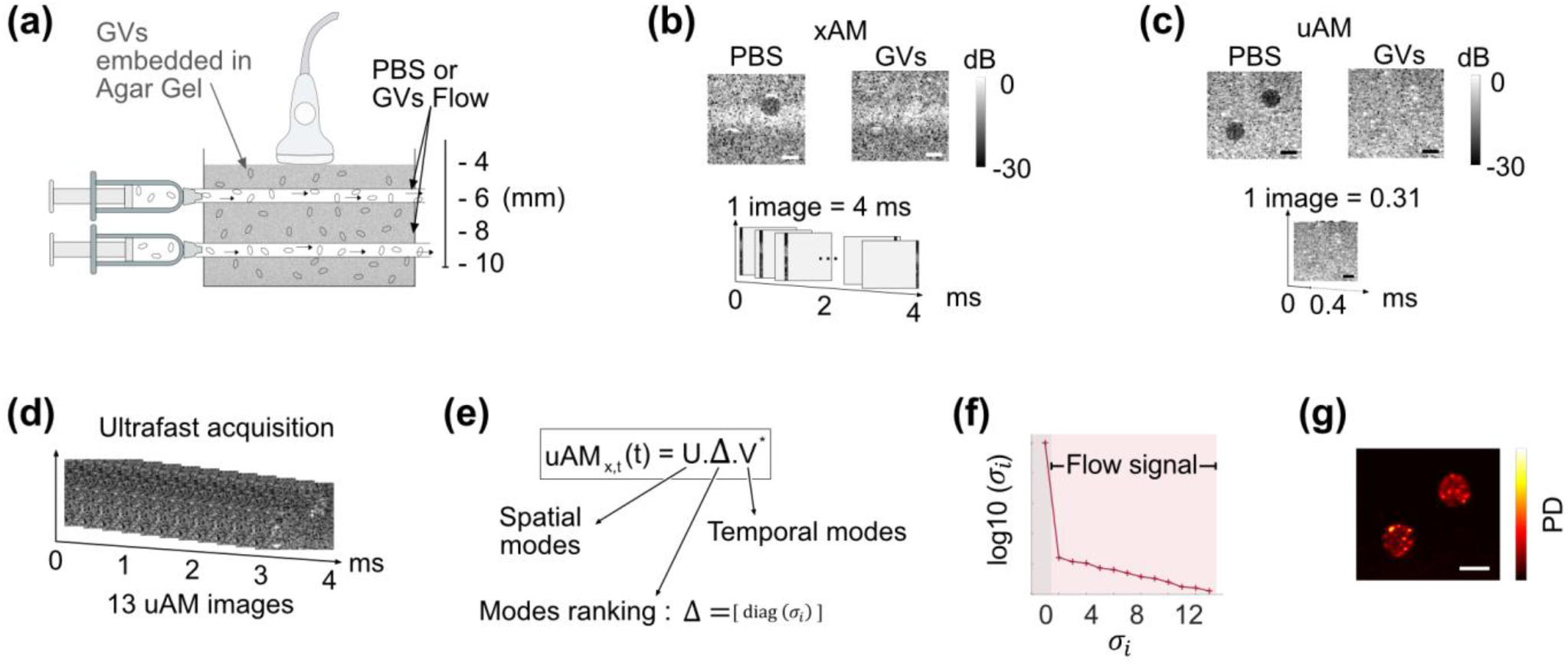
Ultrafast imaging of GV flows in a nonlinear phantom. (a) Illustration of the flow phantom with two horizontal tunnels of diameter 1 mm2 allowing controlled flow of PBS or GVs (114 pM) through an agarose gel phantom containing embedded GVs (114 pM). (b) xAM acquisition of PBS and GV flow conditions. One image of 128 vertical lines is acquired in 4 ms. (c) It takes 0.31 ms to acquire a similar uAM image. (d) In 4ms, 13 uAM images can be acquired. (e) SVD decomposes the 4 ms uAM acquisition of GV flow into a product of separable spatial (U) and temporal (V) matrices weighted by a diagonal matrix Δ. (f) The eigenvalues σi of Δ rank the spatial and temporal modes in the data. A high σi value is associated with stationary background signal whereas the lowest σi values are associated with flow. (g) Excluding the first SVD mode leads to a Power Doppler (PD) image of the GVs flows. Scale bar: 1 mm.

Next, we injected a flow of GVs suspended in PBS at same concentration as in the gel. Now the tunnels blended in with the stationary phantom, making the flowing and stationary GVs indistinguishable in both xAM and uAM. However, in the time it takes to obtain one xAM image, 13 uAM snapshots can be recorded (**Fig.4.d**) and analyzed to create an image specifically of flowing GV using singular value decomposition (SVD)^23^ (**Fig.4.e-g**). This capability to distinguish flowing from stationary nonlinear contrast agents is only possible with the acceleration of uAM.

To examine the utility of uAM in vivo, we leveraged its capacity for simultaneous nonlinear and blood flow imaging to visualize mouse liver function. The removal and degradation of circulating particulates by this organ is critical for homeostasis and response to pathogens, and serves as a diagnostic marker in common diseases^24–27^. GVs can be used as an intravenous contrast agent to visualize and quantify both phagocytic uptake from the blood and lysosomal degradation by liver macrophages^9^. This paradigm tracks the GV-produced enhancement of vascular contrast with power Doppler (PD) imaging and the uptake and lysosomal degradation of GVs in the liver by AM. In the initial study describing this approach, the liver was imaged with xAM, while the vascular contrast had to be measured separately with linear plane wave imaging. We hypothesized that uAM could simultaneously monitor both the molecular and vascular information in the same organ, providing a more practical diagnostic approach.

To test this concept, we injected GVs intravenously in an anesthetized mouse (**Fig.5.a**) and monitored liver PD and AM signals with a single ultrasound probe and uAM sequence over 2800 s, injecting GVs 800 s after beginning the acquisition. Blocks of 200 uAM images (acquired at 500 Hz) were obtained every 8 s. Leveraging the ability of uAM processing to extract simultaneously both linear and nonlinear information from the same pulse sequence (by summing or subtracting the amplitude modulated pulses), we processed two complementary sets of data (**Fig.5.b**, **Appendix**): by applying a clutter filter^23^ to each block of 200 linearly-processed uAM image, we obtained PD images of the liver, tracking the vascular enhancement due to circulating GVs^8^; while the nonlinear-processed images from the same plane tracked the contrast of GVs only. AM images therefore express the population of circulating GVs, but also GVs taken up by liver tissue. The normalized PD and AM signal time courses (**Fig.5.c-d**) show that after injection, the vascular signal quickly reaches a maximum within 60 s, then decreases back to a baseline over 900 s. Meanwhile, the AM signal progressively increases to a maximum around 600 s after the injection, corresponding to the largest concentration of intact GVs in the liver. Macrophages then degrade the GVs, resulting in a gradual decrease towards baseline. The recorded blood and liver pharmacokinetics fit a two-compartment model^9^ whose rate constants parametrize the concurrent processes of phagocytosis and lysosomal degradation. The apparent uptake rate of 0.402 min-1 and degradation rate of 0.0481 min-1 are within the range expected for healthy mice^9^. This in vivo application highlights the capacity of uAM to provide simultaneous access to nonlinear contrast and ultrafast phenomena such as blood flow.

**Figure 5.**
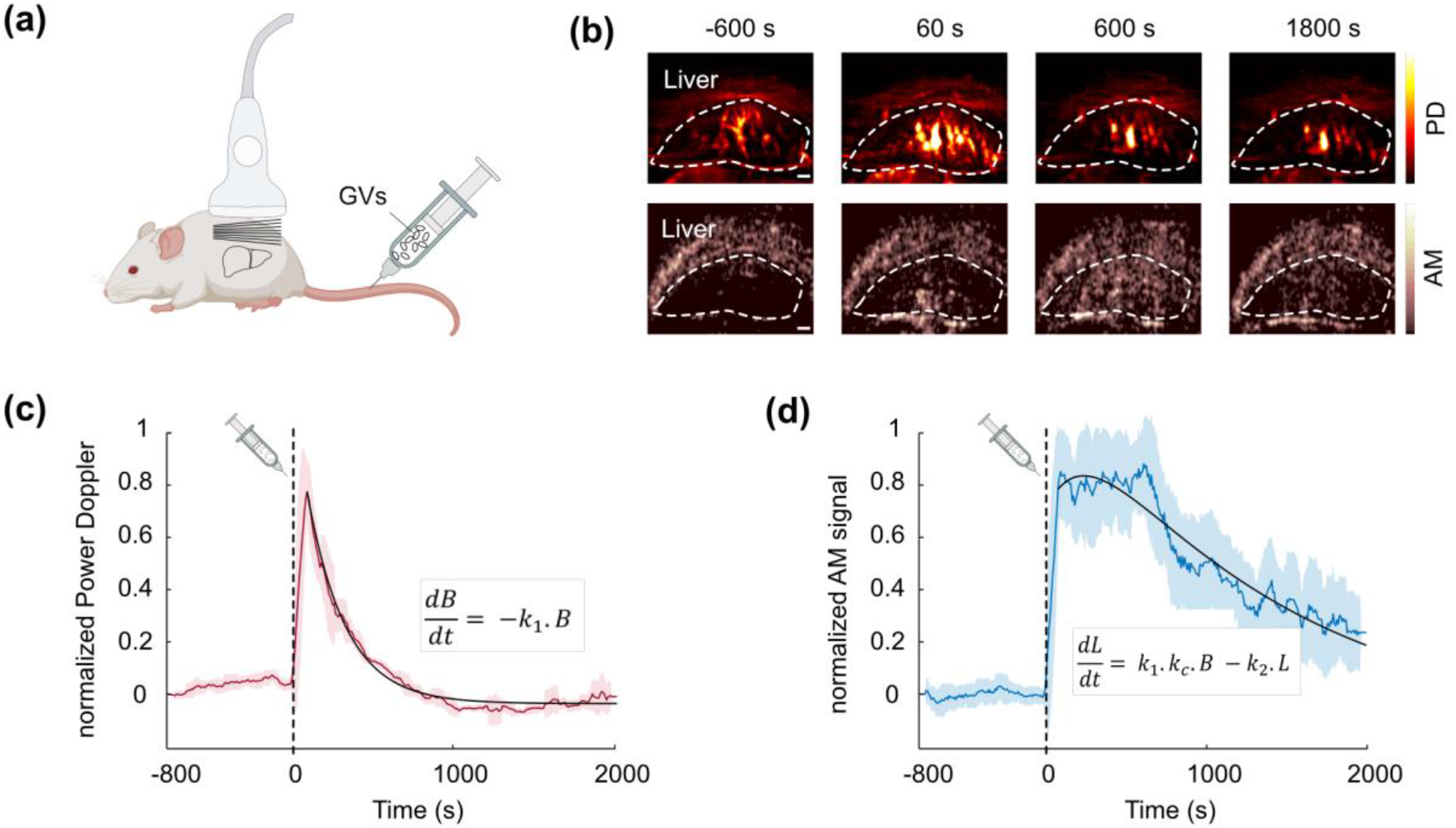
uAM imaging of phagolysosomal function. (a) Schematic representation of the mouse experiment: purified GVs (100 μl, 3 nM) were intravenously injected in an anesthetized mouse, and the liver was continuously monitored with uAM. Blocks of 200 uAM images were acquired every 8 s. (b) Four time points of PD and AM images acquired simultaneously with uAM. The liver is outlined. Scale bar: 1 mm. (c) Normalized liver PD time. (d) Normalized liver AM time course. Dashed black lines: time of GV injection. Continuous colored lines: mean time course averaged over +/-5 neighboring time points (sliding window). Shaded areas: ± STD of neighboring ± 5 time points. Continuous black lines: curves fitted to the data using a pharmacokinetic model where B is the blood signal, L is the liver signal, k1 is the uptake rate, k2 is the degradation rate, kc is a signal scaling factor.

## DISCUSSION

Our results demonstrate that the implemented ultrafast pulse sequence uAM enables the nonlinear imaging of acoustic contrast agents such as GVs with a substantially expanded field of view and dramatically accelerated frame rate compared to existing AM approaches. In addition, it provides simultaneous ultrafast linear acquisition to monitor physiological events such as blood flow. To achieve this remarkable performance, uAM combines amplitude-modulated transmissions with ultrafast MPW imaging and selective coherent compounding of Hadamard-coded echoes. We anticipate that these performance characteristics will make uAM a method of choice for a wide variety of contrast imaging applications. In particular, uAM will facilitate the development of biomolecular contrast agents, reporter genes and biosensors for ultrasound by allowing them track dynamic biological events such as gene expression and enzyme activity across large fields of view and keep up with rapid signaling phenomena. At the same time, the ultrafast linear capabilities of uAM will allow molecular contrast to be visualized within the context of anatomy and physiology, including the blood dynamics imaged in this study, or the motion-tracking of tissue in ultrasound elastography (Genisson et al. 2013).

As with any technique, uAM has some limitations. First, the cumulative propagation of plane waves through nonlinear media can lead to artifacts below nonlinear contrast sources. In scenarios where this is a particular concern, xAM currently provides the best artifact cancellation. Second, MPW transmission creates a small dark zone in the near field of the transducer due to the time required to emit a pulse burst. With 8 angles, this dark zone has a depth of approximately 2mm. In addition, because the half-amplitude plane waves are created in uAM by emitting ultrasound energy with half of the transducer elements, the formation of an acoustic field equivalent of half of the acoustic field created by all the elements requires a longer interference distance after emission. This distance, defined as the near field distance in uAM, was found to be 2.6 mm (at 15MHz, 0.1mm pitch) (Appendix, **Fig.S4**). Finally, potential memory effects in nonlinear objects under successive excitations^22^ were not been taken into account in our study as the measured CAR performance was largely satisfying for our application. If such effects arise in future studies, they could be tackled by implementing orthogonal decoding and pulse-inversion-based harmonic suppression^22^. Despite these limitations, uAM’s exceptional combination of speed, sensitivity and spatial coverage will give this pulse sequence a bright future in contrast ultrasound.

## APPENDIX : Materials and Methods

### uAM transmission sequence

Three ultrasound bursts of modulated amplitude +1, +½, +½, each carrying N successive tilted plane waves are transmitted into the medium one after the other by a linear ultrasound probe with elements along the x-axis, with a reception time separating each burst (**Fig.1.c.i**). These amplitude-modulated multiplane bursts can be written as:

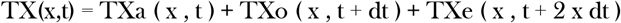

where TXa, TXo, TXe of respective amplitude +1, +½ and +½, each represent the time delay laws of the N successive tilted emissions of different angles [*α_i_*]_*i*=1:*N*_ (**Fig.S1** for N=8). The half amplitude pulses are achieved by silencing the odd (for TXo) or even (for TXe) elements of the ultrasound probe. dt is the time delay between bursts. Within each burst, pulses are emitted one after the other with a delay corresponding to 3 wavelengths between each pulse. *τ_i_* is the time delay of the emission of the pulse *α_i_* after the transmission of the first pulse *α*_1_.

AM radio-frequency (RF) data are generated by subtracting the two backscattered signals elicited by the half-amplitude bursts from the signal elicited by the full amplitude burst. To ensure the largest differential response, the full amplitude burst is set to drive GVs into their nonlinear buckling regime^13^ while the half-amplitude bursts elicit a linear response (**Fig.1.a-b**). We will see later that with the same received data, we can also obtain linear images by summing the contribution of all the received signals (**Fig.1.c.vi**).

The TX triple-transmission is repeated N times (matching the number of angles), with the only difference between the N TX repetitions being the polarization of successive transmitted amplitudes. To obtain an orthonormal basis of TX matrices, the vector [TX TX TX …. TX]_*N*_ is multiplied by the Hadamard Matrix of order N. Hadamard matrices only being of order *N* = 2^*m*^, [m *ϵ* N], the number of angles used in uAM imaging is restricted to values of 2^*m*^. **Fig.1.c-i** illustrates the transmission paradigm of uAM with N = 4 angles. A total of 48 pulses is transmitted in bursts of 4, the burst repeated at 3 different amplitudes and 4 different polarity patterns given by the columns of the Hadamard matrix of order 4.

### Hadamard summation and nonlinear artifacts

In MPW processing, the Hadamard-coded received signals are summed following the combination given by the Hadamard matrix lines. Assuming linear scatters only, the summation of the received signals results in N new ultrasound images, corresponding to backscattered echoes of individual plane waves, but with a virtual amplitude N times greater than that received with a single plane wave.

In AM imaging, the subtraction of the two half-amplitude pulses from the full-amplitude pulse allows the elimination of linear signal and the capture of specifically nonlinear responses. In uAM, these nonlinear signals, produced by trios of modulated MPW bursts, contain Hadamard-coded polarities (**Fig.1.c-ii**). In conventional Hadamard summation, the assumption must be made that that oppositely coded signals sum to zero. However, while this is generally true for linear signals, nonlinear scatterers can produce different echo profiles in response to pulses of opposite polarity that do not cancel each other^18,19^. This phenomenon is the basis for pulse-inversion imaging^21^, and a variety of contrast agents, including GVs, produce pulse-inversion nonlinear contrast^13^.

To illustrate the issue created by pulse inversion nonlinearity, we consider Hadamard summing for a uAM sequence of N = 4. We define p as the AM-subtracted echo from a positive polarity pulse (blue pulse in **Fig.1.c.ii**), n as the AM-subtracted echo from a negative polarity pulse (green pulse in **Fig. 1.c.ii**) and h as the result of summing p + n (black trace in **Fig.1.c.iii**). The four resulting AM RF data can be written as follows (**Fig.1.c.ii**):

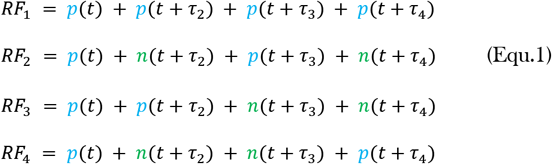

The subtraction and addition of this RF data according to the coefficients of the Hadamard matrix of order 4 then leads to four new sets of Hadamard RF data H_1_, …, H_4_:

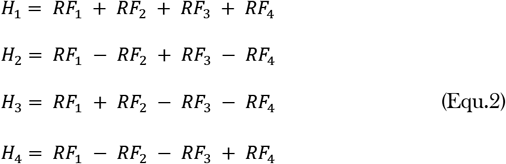

This results in the transformed RF data (**Fig.1.c.iv**):

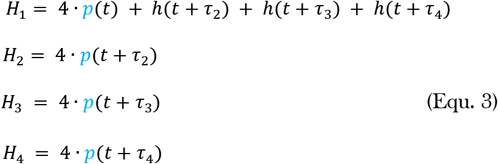

It can be seen from this result that Hadamard subtractions/additions result in the desired positive single-wave signals of quadruple amplitude for H_2_, …, H_4_ but that H_1_ contains both a quadruple-amplitude signal at t and nonlinear signals *h* at (t+τ_2_),(t+τ_2_),(t+τ_2_) due to the pulse inversion operation. This artifactual signal results in the spatial mis-assignment of nonlinear signal during subsequent coherent compounding. This pattern generalizes to higher-order Hadamard matrices. Undesirable nonlinear pulse inversion signals always appear in H_1_, and never in H_>1_. For any MPW sequence of N angles, we have after Hadamard summing of the nonlinear raw data:

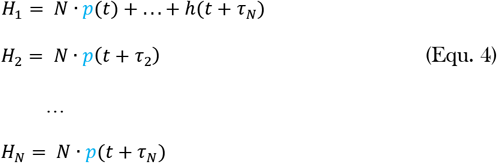

### Selective coherent compounding

Based on Equ. 4 derived above, a selective coherent compounding of all Hadamard RF data except the first one (H_1_) is then performed giving the compounded data set C:

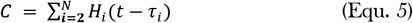

C is then beamformed (delay-and-sum algorithm) and a final nonlinear image S(K_x_, K_z_), (K_x_ and K_z_ being the respective lateral and axial dimensions of the final image) is finally obtained.

### Near field in uAM imaging

In ultrasound imaging, the simultaneous excitation of all the elements of a transducer is assumed to create an acoustic plane wave by interference of the simultaneously-emitted pulses in front of the probe. In order to create two acoustic plane waves of full and half amplitude for uAM imaging, we proposed to emit simultaneously (or with a time delay with respect to each other, to « tilt » the plane waves) ultrasound pulses with all of the 128 elements of the probe (spaced by 0.1 mm), or with every other element (half of them: 64 elements, spaced by 0.2 mm). In the uAM pulse sequence, two half-amplitude fields are created, with the odd and the even elements of the probe. These two half-amplitude fields are then summed to virtually create a full-amplitude acoustic field that is finally subtracted from the field obtained with 128-elements.

We simulated the plane wave formation in front of the probe under the different transmission conditions of uAM (all or half of the elements emitting) using k-wave (open-source acoustics toolbox for MATLAB). We simulated wave propagation in a homogeneous and isotropic medium (water): the speed of sound was set to 1480 m/s. The size of the domain was 12.8 mm × 12.8 mm and was discretized with a step size of 50 μm; perfectly matched layers were used to absorb the waves at the edges of the domain; 128 transducer elements were positioned 1 mm from the top of the domain and positioned horizontally 0.1 mm from each other. The source broadcasted a short pulse (1 cycle) with a central frequency of 15 MHz. **Fig.S4.a** shows the propagation of the acoustic field after simultaneous emission by all of the 128 elements (Pressure field: Pa), while **Fig.S4.b** shows the propagation of the acoustic field after simultaneous emission by 64 elements only (the odd elements, pressure field: Po), spaced by 0.2 mm. In a third simulation (not shown), we calculated the resulting acoustic field obtained after emission by the 64 even elements of the transducer (pressure field: Pe). We then compared the resulting maximum pressure fields at different depths obtained with Pa or with the sum of Po+Pe (**Fig.S4.c**). The resulting wave in front of the probe is mostly planar between the lateral coordinates [+1mm; +11.8mm], outside of the edge effect regions on the array’s extremes. In this analysis, the mean value of the maximal received pressure at a given depth and the standard deviation of the horizontal pressure fields is calculated between these coordinates.

In the near field (which will be defined by the end of this section), the sum of the two half-amplitude fields don’t equal the full-amplitude field. At 1 mm depth, for example (**Fig.S4.c**), the maximum pressure field is substantially higher for Po+ Pe than for Pa. This observation is confirmed **Fig.S4.d**, which shows that the mean Po+ Pe maximal pressure at depth 1 mm is 723 kPa whereas the maximal Pa pressure is averaged around 618 kPa. However, at greater depth, the averaged field Po+ Pe gets closer to the mean value of the field Pa. At a depth of 2.7 mm, Po+ Pe = 638 kPa and Pa = 636 kPa, the difference being below 0.4%. With less than 0.5% difference, we consider the two fields as substantially equal.

We also considered the standard deviation of the pressure field (along the horizontal axis) as a function of the depth. At each depth, we evaluated the ratio of the standard deviation and the mean pressure value Po+ Pe and Pa (**Fig.S4.e**). Close to the probe (< 2.5 mm depth), the standard deviation ratio of the pressure field is higher for Po+ Pe (>4%) than for Pa (<2%). At 2.6 mm, this ratio for Po+ Pe drops to 1.7% and equates to the ratio of Pa within an accuracy of 0.03%.

We define the near field in uAM imaging as the depth for which the pressure fields created by all (128), or by the sum of the odd (64) and the even (64) elements generate equal maximum peak values (averaged along the horizontal line defined by the depth) within an accuracy of 0.5% and for which the standard deviations (for each transmission condition) of the maximum pressure field at this depth represent less than 2% of the average pressure. At 15 MHz, with a 0.1 mm pitch ultrasound probe, the near field in uAM is therefore 2.6 mm.

### Spatial resolution calculation

The ultrasound phantom for quantifying the resolution of the imaging techniques was prepared by embedding GVs in 1% (w/v) agarose gel in PBS. GVs were incubated at 42°C and mixed in a 1:1 ratio with molten agarose for a final GV concentration of 1 OD500nm and loaded into a custom rectangular mold. To make the pattern of repeated lines of intact GVs (**Fig.S3.a**), a 3.7MHz acoustic standing wave was applied to the phantom using a custom device such that the peak positive pressure at the pressure antinodes of the standing wave exceeded the critical collapse pressure of the GVs^28^. The driving pressure was selected such that the line pattern generated by the remaining intact GVs was less than 100 μm in width. The ground truth of the patterned GVs was imaged using brightfield optical imaging (**Fig.S3.b**), and quantified using ImageJ, the full width at half-maximum (averaged across N=13 lines) was measured as 60 μm. We positioned the phantom in a water bath with the lines being oriented horizontally, and imaged it successively with pAM, xAM and uAM (**Fig.S3.c**). We repeated the imaging protocol by flipping the phantom of 90º and imaged the vertical lines (**Fig.S3.d**). The spatial axial and lateral resolutions of each pulse sequence were calculated by evaluating the full width at half-maximum of the horizontal (axial resolution) and vertical (lateral resolution) lines (averaged over 20 lines and 10 time points).

### Linear processing

All the received RF data are stored in the ultrasound scanner, allowing multiple different signal processing operations in addition to nonlinear imaging. For linear imaging, we sum the full and half-amplitude pulses instead of subtracting them (**Fig.1.c.vi**) and repeat the remaining post-acquisition processing described in the previous section. This results in B-mode ultrasound images, free from nonlinear artifacts thanks to selective coherent compounding.

### Dynamic processing of ultrafast images

Ultrasound plane wave imaging enables ultrafast scanning of the media16. The ultrafast repetition of the uAM pulse sequence therefore provides access to fast transient linear and nonlinear events in the whole imaging plane. In two example applications of this capability, we applied a clutter filter to the spatiotemporal data^23^ to detect fast flows in the imaging planes.

### Ultrasound acquisition sequence

Ultrasound sequences were implemented on a research ultrasound scanner (Verasonics, USA) using an ultrasound probe with 128 linear elements emitting at 15.625 MHz (pitch = 0.10 mm). The acquisition scripts and processing codes were written in Matlab (Matworks, USA).

We used a Verasonics Vantage ultrasound system with an L22-14v probe (Verasonics Inc., Redmond, WA, USA) to implement the uAM, xAM and pAM imaging sequences. The probe is a linear array of 128 elements with a 0.10-mm pitch, an 8-mm elevation focus, a 1.5-mm elevation aperture, and a center frequency of 18.5 MHz with 67%–6 dB bandwidth. We applied a single-cycle transmit waveform at 15.625 MHz to each active array element to ensure our fundamental frequency is divided 4 times with the 62.5-MHz sampling rate of the system. For the uAM sequence, we use the full aperture of the probe, i.e., 128 elements, to send the successive plane waves. For xAM, to reproduce the optimal results published in Maresca et al.15, we used an aperture of 65 elements for the xAM sequence. For pAM, we used an aperture of 89 elements. The input voltage driving the transducer was set in order to reach, for each modality, a peak positive pressure of 500kPa.

### Framerate evaluation for pAM, xAM and uAM

For a 10mm depth acquisition, theoretically acquired along the 128 elements of a transducer linear array:

One xAM image is acquired after the transmission along each vertical line of an image of three distinct pulses. In xAM these three pulses correspond to the two individual and one cross propagating plane wave.

One image is then acquired in:

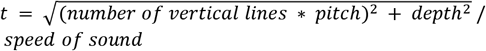

with: number of vertical lines = 128, pitch = 0.10 mm, depth = 10 mm and speed of sound = 1540 m/s

so: 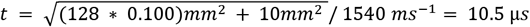

Then, one image of 128 vertical lines can theoretically be acquired in: *T_image_*(*xAM*) = 10.5 μ*s* × 3 × 128 = 4.03 *ms* which corresponds to an imaging framerate of 250Hz.

The same calculation can be applied for pAM imaging and would result in the same imaging framerate.

One similar uAM image of 128 vertical lines is acquired after the transmission of 3 amplitude modulated sets of 8 multiplane wave bursts. After the emission of the first tilted angle, 2.4 additional microseconds are necessary to transmit the rest of the 8-multiplane wave burst (see **Fig.S1**). A uAM transmission is therefore 2.4 μs longer than a single plane wave transmission. Therefore, the backscattered echoes from the last plane wave of the burst arrive with a delay of 2.4 μs after the echoes coming from the first plane wave of the burst. Then, the uAM transmission/reception period is: 10.5 + 2.4 = 12.9 μs. One uAM-8 image therefore can be acquired in: *T_uAM_* = 12.9 μs × 3×8=0.309 ms which corresponds to an imaging framerate of 3.2kHz.

For the same field of view (width given by the probe: 128 vertical lines of 0.1mm, depth: 10mm), uAM is therefore 3.2/0.250 = 12.8 times faster than conventional xAM or pAM imaging.

These calculations are performed to compare imaging framerates of same-width images acquired by the different techniques. In reality, the imaging aperture used in in our experiments spanned 89 elements of the probe for pAM15 and 64 elements for in xAM^15^. The imaging framerate for these narrower images would therefore be proportionally accelerated, but at the cost of smaller field of view. Therefore in practice, as implemented in this study, the number of transmits in uAM is reduced by more than 8-fold compared to pAM or xAM for the formation of one AM image

### Preparation of gas vesicles

Nonlinear GVs were prepared as described in Lakshmanan et al.^29^. Briefly, GVs were harvested from buoyant Anabaena flos-aquae cells by hypertonic lysis and purified by repeated centrifugally assisted flotation and resuspension. The GVs were stripped of their outer GvpC layer by treatment with 6M urea, followed by additional repeated centrifugally assisted flotation, dialysis in 1x PBS and resuspension to remove the GvpC and urea.

### Tissue-mimicking phantoms

Tissue-mimicking phantoms for imaging were prepared by casting 1% (w/v) agarose in PBS with 0.2% (w/v) AlO3. For static imaging, we used a custom 3D-printed mold to create two rows of cylindrical wells with 2 mm diameter. GVs were incubated at 42 °C for 1 minute and then mixed in a 1:1 ratio with low-melt molten agarose (at 42 °C) for a final GV concentration corresponding to 3 OD500nm and loaded into the phantom. The AlO3 concentration was chosen to match the scattering echogenicity of the GV well. According to 30, the molarity of GVs is 114 pM/OD. The upper row was centered at 6 mm depth and the lower row at 9 mm.

For dynamic imaging, using a custom 3D-printed mold through which 1 mm diameter tubes are arranged, we mixed in a 1:1 ratio of GVs (OD 2) and molten agarose for a final GV concentration corresponding to 1 OD500 nm and loaded into the phantom. After 20 min, we made sure that the agarose was fully solidified and gently pulled out the tubes leaving 1 mm diameter tunnels crossing the whole agarose phantom. Solutions of PBS or of GVs in suspension (OD 1) were then injected through the tube using a syringe pump at 1 mL/min.

Extracting signal from flowing GVs only during the dynamic in vitro study

To distinguish the flow of GVs from the stationary ones in the agarose phantom in 4ms only with uAM imaging, we performed a SVD decomposition on the 13 uAM-nonlinear images in order to decompose the acquisition into a weighted, ordered sum of separable spatio-temporal modes expressed by U, Δ and V as:

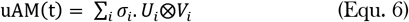

with U_i_ and V_i_ the i-th columns of the corresponding SVD matrices U and V. And Δ the weighting matrix. U represent the spatial-only modes of the uAM acquisition and V the temporal-only modes. Finally, the diagonal matrix Δ ranks these modes. As described in^23^, removing the first mode(s) from the SVD decomposition allows the extraction of the flow-only from the acquisition. Here, we remove the first mode only and apply an inverse SVD decomposition to reconstruct a 3D (2D space + 1D temporal) acquisition without the background signal component. The PD value PD for each pixel is then calculated as:

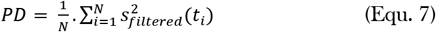

Where N=number of temporal samples in one uAM-doppler block (here N=13) and s_filtered_ is the filtered signal.

### In vivo imaging

All animal experiments were conducted under protocols approved by the Institutional Animal Care and Use Committee of the California Institute of Technology. The in vivo experiment presented has been performed on a BALB/c female mouse (Jackson Laboratory) aged 7 weeks under a protocol approved by the Institutional Animal Care and Use Committee of the California Institute of Technology. No randomization or blinding are necessary in this study. The mouse was anesthetized with 2%–3% isoflurane, depilated over the imaged region, and imaged using an L22-14v transducer with the uAM pulse sequence. Eight hundred seconds after the start of imaging, 120 μL of OD500 25 GVs were infused over 10s. The acquisition continued for 2000 s after injection.

### In vivo PD processing from uAM-doppler acquisitions

Power Doppler visualizes the concentration of moving blood cells during a determined time interval ΔT. This integration time should cover several cardiac cycles in order to average the heartbeat dynamic. The averaged heartbeat frequency in mice is estimated at 10 Hz. A time interval ΔT = 400 ms is therefore adequate to average approximately four heartbeats. uAM images were acquired at 500 Hz, which is equivalent to 200 images during 400 ms, with one uAM-doppler block repeated every 8 s. To extract the signal from the blood stream only (in which both red blood cells and GVs contribute to the signal), linear processing is performed on the received AM data after transmission of the uAM pulse sequence: instead of subtracting the half-amplitude pulses from the full-amplitude pulses for nonlinear processing, the summation of the AM backscattered echoes is performed to conserve the linear components of the ultrasound image (**Fig.1.c.vi**). A clutter filter as described in the dynamic in vitro study and in ^23^ is then applied to each of the uAM linearly-processed blocks (of 200 images acquired at 500Hz) to extract the blood signal from the spatiotemporal acquisition. For this study, the first 50 SVD modes were removed from the SVD decomposition. The PD value PD for each pixel is then calculated as:

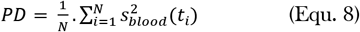

Where N=number of temporal samples in one uAM-linear doppler block (here N=200) and s_blood_ is the filtered blood signal.

A similar processing scheme could be applied to nonlinear uAM-nonlinear doppler images (images obtained after subtraction of the half-amplitude pulses from the full amplitude ones to conserve the nonlinear components of the image only) and will therefore express the PD of moving GVs only (S2):

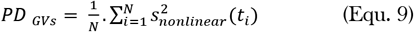

at 3.75 MHz, 7.6 V peak-to-peak. Images of the channel were acquired for 10 seconds during ultrasound application as described above.

## ACKNOWLEDGEMENTS

This research is supported by the National Institutes of Health (R01-EB018975). C.R. is supported by the Human Frontier Science Program (Grant No. LT000217/2020-C). Related research in the Shapiro laboratory is supported by the Chan Zuckerberg Initiative, the Pew Charitable Trust, the David and Lucile Packard Foundation and the Heritage Medical Research Institute.

## AUTHOR CONTRIBUTIONS

C.R. and M.G.S. conceived the study. C.R. developed sequence acquisitions and acquired data. C.R. and D.W. developed the processing model. D.M. produced the GVs. Z.J. helped conceiving the dynamic in vitro study. B.L. assisted with the in vivo study. C.R. and M.G.S wrote the first draft of the manuscript. All authors edited and approved the final version of the manuscript.

