## Supplementary figures for "Ultrafast amplitude modulation for molecular and hemodynamic ultrasound imaging"

**%%%%%%%%%%%%%%%**

**Supplementary Figures**

**%%%%%%%%%%%%%%%**

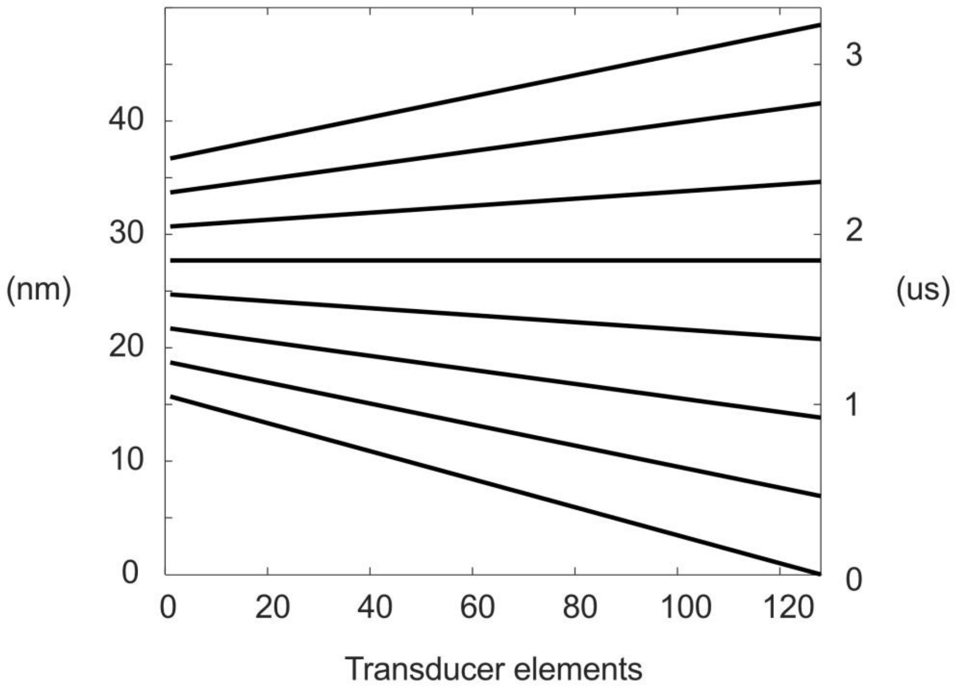


Figure S1: Time delay law of one 8-angle burst of the uAM transmission sequence.

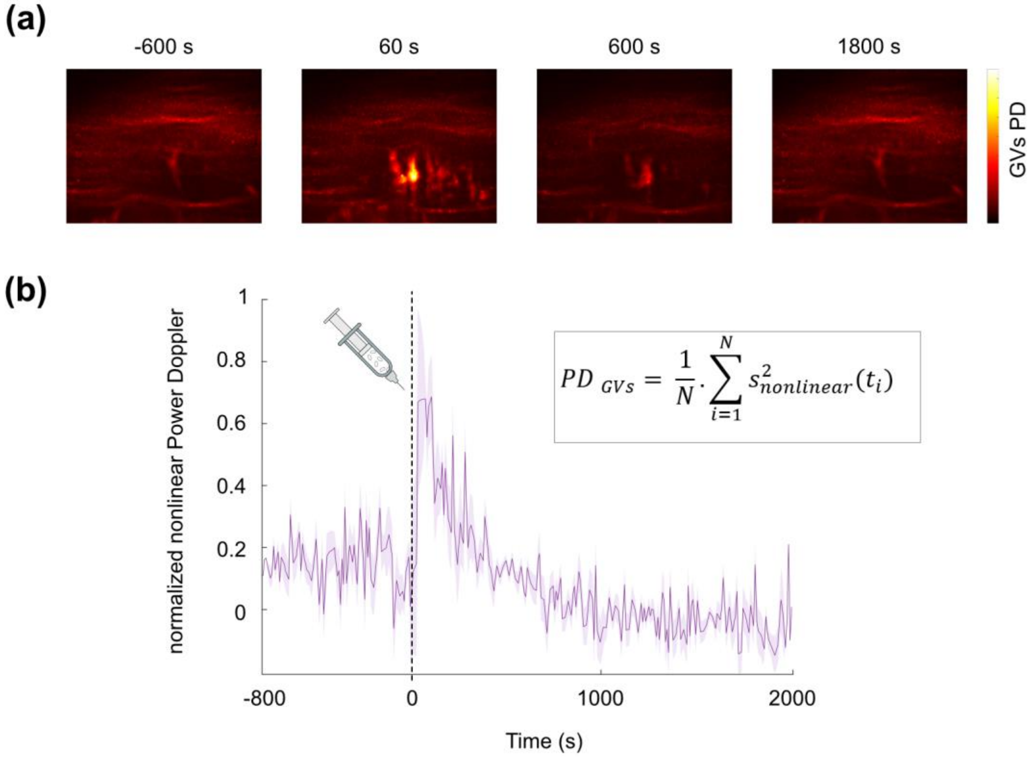


**Figure S2: Power doppler processing of the nonlinear signal during mouse liver imaging.** (a) The power Doppler images illustrates the absence of circulating GVs before injection (-600s), the uptake of GVs right after injection (60s) and clearing of GVs from the blood stream despite the presence of GVs in the liver (>600s). (b) Nonlinear PD time course within the liver.

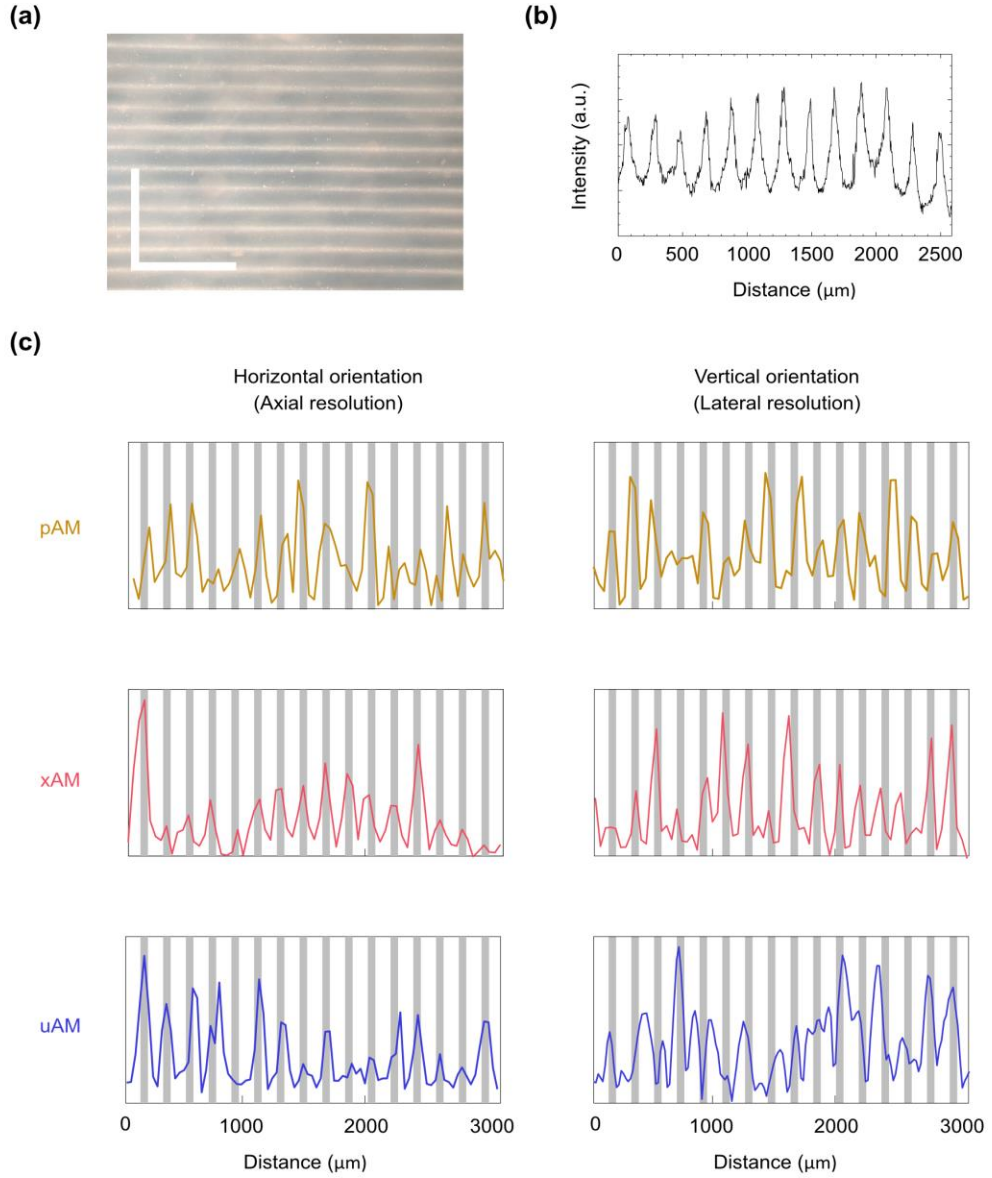


**Figure S3: Spatial resolution estimation of pAM, xAM and uAM imaging using an agar phantom with repeated lines of intact GVs**. (a) Bright field optical imaging of the agar phantom. White lines correspond to GVs. Scale bar: 1 μm (b) Optical intensity profile measured along a single line of the phantom. The ground truth of the patterned GVs was quantified using ImageJ: the full width at half-maximum was measured as 60 μm (c) pAM, xAM and uAM intensity profile of the phantom in the horizontal, and vertical orientation.

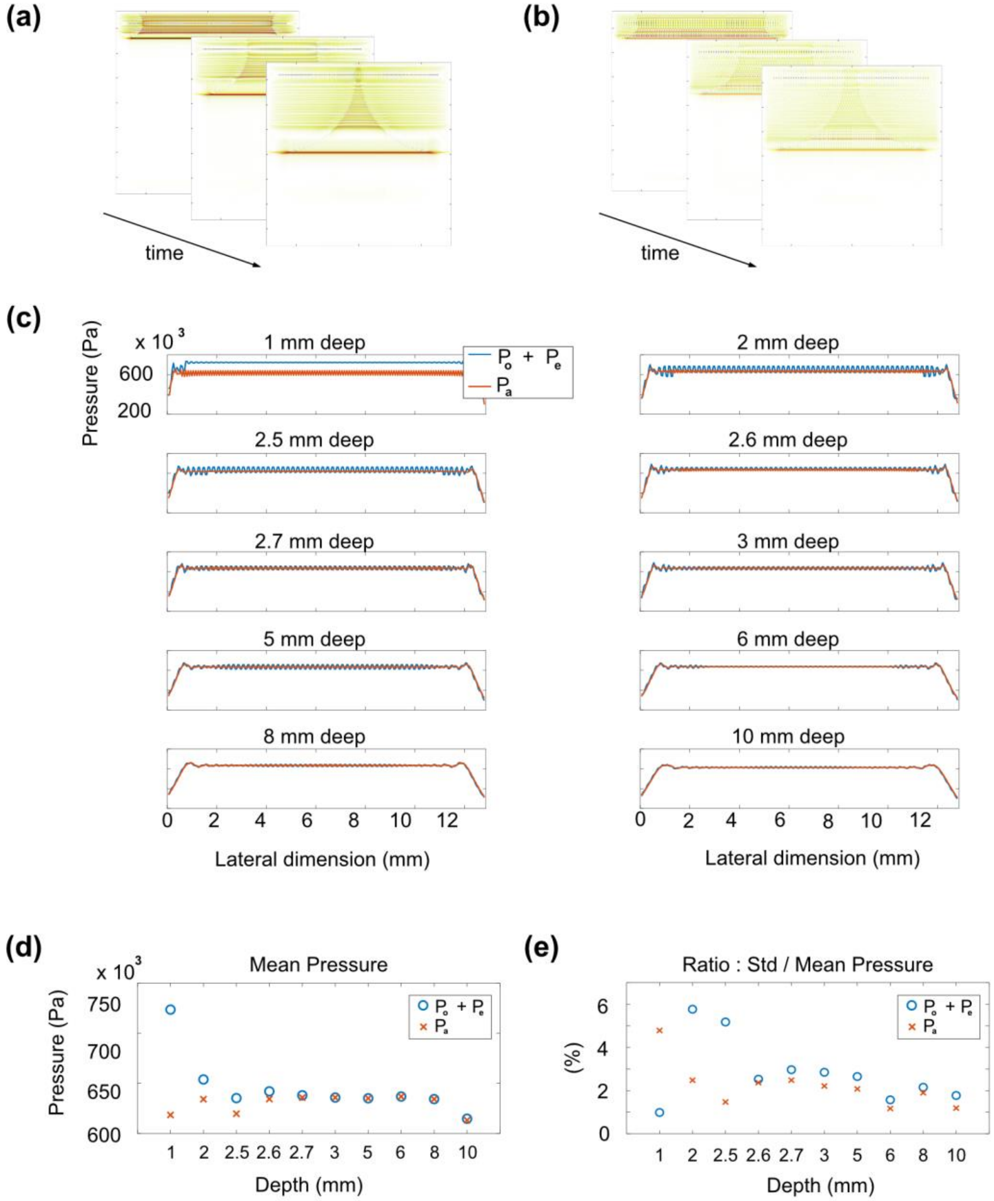


**Figure S4: Acoustic pressure field k-wave simulation of plane waves generated with 128 or 64 elements**. (a) k-wave spatio-temporal simulation of the acoustic field Pa created by the simultaneous emission of the 128 transducer elements of an ultrasound probe (15MHz pulse, 1 cycle, pitch 0.1 mm). (b) Spatio-temporal simulation of the acoustic field created by the simultaneous emission of 64 elements of the same ultrasound probe (Po : odd elements only, now spaced by 0.2 mm). (c) Lateral pressure profiles of the maximum received pressures at different depths: Pa corresponds to the pressure propagating after emission by the 128 elements of the probe. Po+ Pe corresponds to the sum of the pressures from the odd (64 elements) and even (64 elements) elements of the probe. (d) Mean lateral pressure at different depths calculated between the lateral coordinates +1 mm and +11.8 mm of the probe e. Ratio of the standard deviation (Std) and the mean lateral pressure calculated between +1 mm and +11.8 mm
